# The lateral habenula mediates the association of a conditioned stimulus with the absence of an appetitive unconditioned stimulus

**DOI:** 10.1101/2022.04.27.489815

**Authors:** Dong-Hee Kim, In-Beom Jin, Nam-Heon Kim, Yong-Jae Jeon, Bo-Ryoung Choi, Michela Gallagher, Jung-Soo Han

## Abstract

The lateral habenula (LHb) has been implicated in conditioned inhibition. Here, the modulating effects of LHb activity on the association of a conditioned stimulus (CS) with the nonoccurrence of an unconditioned stimulus (US) were assessed in vivo using chemogenetic methods. Animals initially received explicitly unpaired CS light and US food presentations. Animals subjected to a retardation-of-acquisition task subsequently underwent light and food pairings, whereas those subjected to a summation task underwent compound light-tone and food pairings. The inhibitory light strength was assessed based on retardation of the acquisition of food-cup conditioned responses (CRs) in light-food pairings and comparisons of food-cup CRs to each stimulus in a CS-alone test following compound training. Neurotoxic LHb lesions and chemogenetic LHb inhibition throughout unpaired training attenuated the inhibitory light strength. Furthermore, chemogenetic LHb activation accelerated the decline in CR induced by repeated light-alone presentations following light-food pairings. Therefore, the LHb critically contributes to conditioned inhibition.

## Introduction

Exposure to a contingency between a neutral stimulus or conditioned stimulus (CS) and a biologically significant stimulus or unconditioned stimulus (US) modifies the behaviors of an organism to the CS (Pavlov, 1927; Mackintosh, 1983; Rescorla, 1988). Such experiences can establish an excitatory association between the two events that elicits a learned response, such as approaching a food-cup area (Holland, 1993; Holland and Gallagher, 2004; Holland and Schiffino, 2016). In contrast, conditioned inhibition is mediated by a CS association with the nonoccurrence of a US that inhibits the operation of conditioned excitation, in this example manifesting as not approaching a food-cup area (Pavlov, 1927; Rescorla, 1968, 1969a; Wasserman et al., 1974; Hearst and Franklin, 1977). Pavlovian conditioned excitation and inhibition are both fundamental mechanisms that guide adaptive behavior (Holland, 1984; Sosa and Ramirez, 2019).

Research in this area, in which a focus on conditioned excitation has long dominated, has revealed that neural substrates of appetitive and aversive conditioned excitation are the amygdala and its interconnected neural structures (Holland, 1984; Gallagher and Holland, 1994; Johansen et al., 2011; Holland and Schiffino, 2016). In addition, the nature of conditioned inhibition has been defined and elucidated in a study on conditioning (Sosa and Ramirez, 2019). However, its neural substrate has remained poorly understood (Sosa et al., 2021), although recent studies have implicated that the lateral habenula (LHb) plays a role in responding to a CS-US contingency in appetitive and aversive learning (Matsumoto and Hikosaka, 2007; Stamatakis and Stuber, 2012; Trusel et al., 2019).

The LHb, an evolutionally conserved region, receives inputs from forebrain limbic structures critical for appetitive and aversive conditioned excitation and provides an inhibitory influence on midbrain dopaminergic neurons (Christoph et al., 1986; Kim and Lee, 2012; Brown and Shepard, 2016; Mizumori and Baker, 2017). Midbrain dopamine neurons are activated by a cue signaling the presence of a reward and are depressed by a cue signaling the absence of a reward (Schultz, 1998; Tobler et al., 2003). Conversely, LHb neurons are activated by a cue predicting the omission of a reward and are suppressed by a cue signaling the presence of a reward (Matsumoto and Hikosaka, 2007). Our recent study measured c-Fos levels in the LHb, substantia nigra pars compacta (SNc), and ventral tegmental area (VTA) of rats that underwent training in which a visual CS that was either paired (positive contingency) or explicitly unpaired (negative contingency yielding conditioned inhibition) with food (Rescorla, 1969b; Matzel et al., 1988; Williams and Overmier, 1988; Droungas and LoLordo, 1995). The levels of c-Fos in the LHb were higher in unpaired animals than in paired animals, whereas c-Fos levels in the SNc/VTA were higher in paired animals than in unpaired animals (Choi et al., 2020). Furthermore, LHb lesions disrupt outcome-specific conditioned inhibition generated by backward conditioning, in which the US precedes the CS (Laurent et al., 2017), although learning through backward conditioning has been reported to be excitatory, inhibitory, or both, depending on the testing circumstance (Barnet and Miller, 1996; Cole and Miller, 1999; Urushihara, 2004).

Therefore, we hypothesized that the LHb would mediate the association of a CS with the absence of an appetitive US. Here, we used designer receptors exclusively activated by designer drugs (DREADDs) to assess the modulating effects of LHb activity on the acquisition of CS-no US contingency learning using retardation-of-acquisition and summation task procedures for measuring conditioned inhibition (Rescorla, 1969a). Animals subjected to retardation-of-acquisition and summation tasks first received explicitly unpaired training (light-nonoccurrence of food) to generate conditioned inhibition. In the second phase, the animals underwent light and food pairings for the retardation-of-acquisition task and compound light-tone and food pairings for the summation task. For the summation task, food-cup responses to either light- or tone-alone presentations were measured in the test phase. The strength of light as a conditioned inhibitor was assessed based on retardation in the acquisition of light-food excitatory conditioned responses (CRs) and comparing food-cup responses to each CS. Experimental LHb manipulations, namely LHb lesions and chemogenetic LHb inhibition throughout the unpaired training, attenuated the inhibitory strength of the explicitly unpaired light. Extinction also generates conditioned inhibition (Calton et al., 1996). Our recent study reported that c-Fos levels were increased in the LHb of rats that received repeated light-alone presentations following light-food pairings and that neurotoxic LHb lesions decelerated the extinction of food-cup CRs (Kim et al., 2021). Accordingly, we predicted that chemogenetic LHb activation would accelerate the extinction of food-cup CRs. We found that chemogenetic activation of the LHb during repeated CS-alone presentations facilitated a decline in food-cup CRs. All results of the present study support our hypothesis that the LHb is a crucial brain region for assessing the association of a CS with the nonoccurrence of a US.

## Results

### Neurotoxic LHb lesions prevent decremental effects of unpaired learning on subsequent paired learning

Recently, we examined the effects of LHb lesions on the acquisition of paired and unpaired learning and found no differences in the acquisition of conditioned behavior between sham-operated and lesioned rats throughout the 8-day paired and unpaired training protocol (Choi et al., 2020). Therefore, we assessed the inhibitory properties of the explicitly unpaired CS by joint application of retardation and summation tasks, as recommended by Rescorla (Rescorla, 1969b), and examined the effects of experimental LHb manipulations on an inhibitory CS. The association of a CS with the nonoccurrence of a US generates conditioned inhibition (Rescorla, 1969b; Matzel et al., 1988; Williams and Overmier, 1988; Droungas and LoLordo, 1995). Because the CS controls a response tendency towards opposing excitation during unpaired training (Rescorla, 1969b), we first used the retardation-of-acquisition procedure, which measures the inhibitory effect of a CS (Figure 1A). The premise of this procedure is that the prior association of a CS with the nonoccurrence of a US retards the acquisition of subsequent excitatory learning. Therefore, according to our hypothesis, no retardation would occur in rats with LHb lesions (Figure 1B).

**Figure 1.**
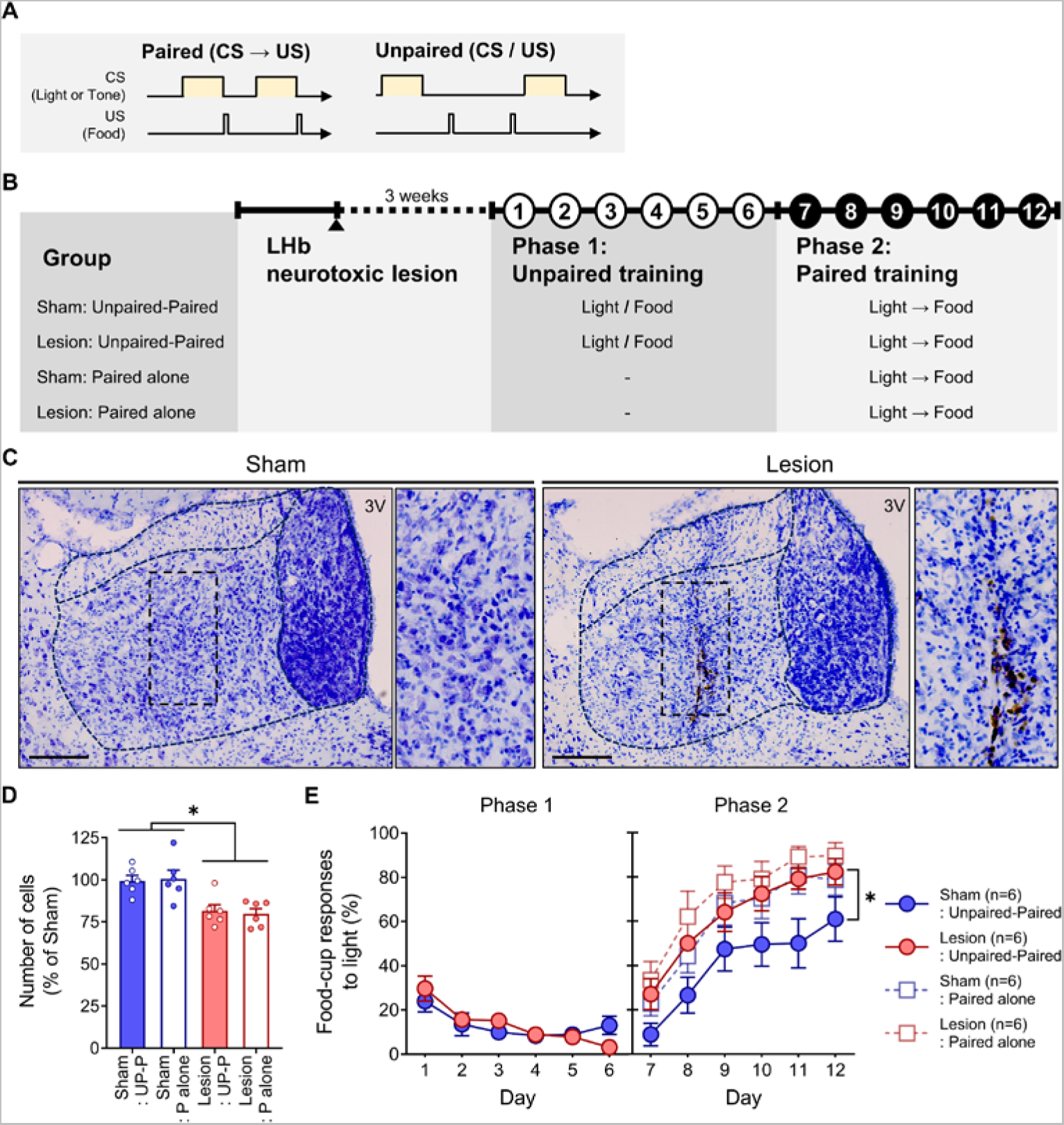
Neurotoxic LHb lesions prevent decremental effects of unpaired learning on subsequent paired learning. **(A)** Conditioning procedures. **(B)** Experimental schedule. **(C)** Photomicrographs showing Nissl-stained control (left) and neurotoxin-lesioned LHb (right), with evident loss of neurons. Magnified images of the boxed area denoted by the dotted line are presented at right. Scale bar, 200 µm. 3 V, third ventricle. **(D)** Neural densities were significantly decreased (to ∼25%) in the lesioned group compared with the sham group (*). **(E)** No effects of LHb lesion were observed in unpaired training (left), but were noted in paired training (*), showing that the acquisition of conditioned food-cup responses was retarded in sham-operated rats with prior unpaired training, but not in lesioned rats with prior unpaired training (right). Error bars represent SEM.

Figure 1C shows photomicrographs of Nissl-stained sections of sham-operated and lesioned rats. Control rats exhibited regular-sized, granule-shaped LHb neurons. In contrast, LHb neurons in lesioned rats were small and kite-shaped. To quantify the degree of damage, we measured neuronal densities in the LHb (Figure 1D). A two-way analysis of variance (ANOVA) of cell densities with training and lesion revealed a significant lesion effect (F_(1,20)_ = 25.27, *p* < 0.001) with no training effect (F_(1,20)_ = 0.004, *p* = 0.95) or lesion × training interaction effects (F_(1,20)_ = 0.15, *p* = 0.71). The left panel in Figure 1e shows food-cup responses to light during unpaired training in phase 1. A two-way repeated-measures ANOVA of conditioned food-cup responses with lesion and day as the main factors revealed a significant effect of day (F_(5,50)_ = 9.07, *p* < 0.001), but no effect of lesion (F_(1,10)_ = 0.37, *p* = 0.85) or lesion × day interaction (F_(5,50)_ = 1.42, *p* = 0.23), indicating that food-cup responses to light during unpaired training were significantly decreased in sham-operated and lesioned rats.

The right panel in Figure 1E shows food-cup responses to light in the paired training of sham-operated and lesioned rats, with or without prior unpaired training, in phase 2. A three-way repeated-measures ANOVA with lesion, training (prior vs. no prior unpaired training), and day as the main factors revealed a significant effect of day (F_(5,100)_ = 9.07, *p* < 0.001), lesion (F_(1,20)_ = 7.38, *p* < 0.05), and training (F_(1,20)_ = 5.98, *p* < 0.05), but showed no interaction effects among the main factors. All four groups exhibited increased food-cup responses to light during paired training. Figure 1E shows that sham-operated rats with prior unpaired learning presented lower levels of food-cup responses to light during paired training, but lesioned rats with prior unpaired training presented levels of food-cup responses to light on par with those of sham-operated rats and lesioned rats with no prior training. Lesioned rats exhibited no decremental effects of unpaired training on subsequent paired learning. The frequencies of food-cup responses were also measured 5 seconds before the presentation of CS in all experiments, including the lesion experiment. No differences between groups were observed (Supplement table 1).

Because preexposure to a CS-alone can also retard the acquisition of a subsequent excitatory association, a phenomenon termed latent inhibition_33_, we also estimated the contribution of this alternative account to the retardation. Preexposure to the same number of CS presented in the unpaired training did not significantly retard the acquisition of subsequent excitatory association, and there was no LHb lesion effect (Figure 1-Supplement figure 1).

### Chemogenetic LHb inhibition throughout unpaired learning blocks the decremental effects of unpaired learning on subsequent paired learning

LHb neuronal activities were inhibited by micro-infusing an adeno-associated virus (AAV) vector into the LHb to express human Gi-coupled M4 muscarinic receptor (hM4Di)-DREADDs under the control of the human synapsin 1 promoter. The inhibitory DREADD, hM4Di, has been experimentally evolved to respond to the synthetic agonist, clozapine-*N*-oxide (CNO). Successful infection of the LHb area was confirmed based on the localized expression of mCherry (53.38% ± 3.50%) or enhanced green fluorescent protein (EGFP; 58.78% ± 8.79%) (Figure 2B–E). Vehicle (VEH)-treated (Gi+VEH:Unpaired-Paired) and CNO-treated (Gi+CNO:Unpaired-Paired) groups were designed to examine the effects of reduced LHb activity on retardation of acquisition. A control group (EGFP+CNO:Unpaired-Paired) was added to exclude the effects of CNO or transfected gene (EGFP) on retardation of acquisition (details in the Methods, Figure 2A). An additional Gi+VEH;Paired alone group was also included for comparison.

**Figure 2.**
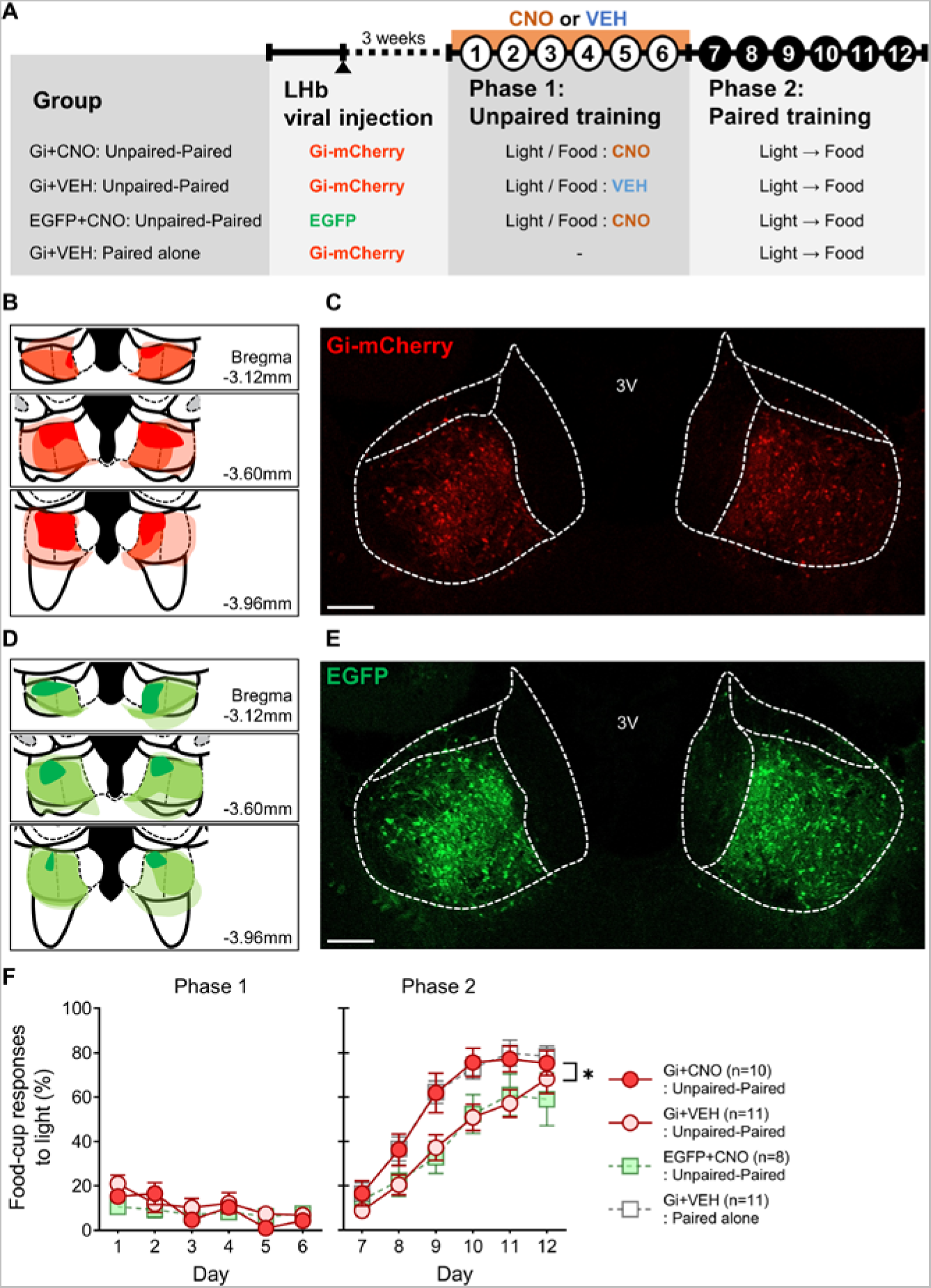
Gi-DREADD–mediated inhibition of the LHb throughout explicitly unpaired training blocks retardation of acquisition. **(A)** Experimental schedule. **(B, D)** Histological reconstruction showing the largest (bright), average (medium), and smallest infected areas (dark). **(C, E)** Representative photomicrographs of the LHb expressing virally transduced hM4Di mCherry or EGFP. Scale bar, 200 µm. **(F)** Gi+VEH and EGFP+CNO rats with prior unpaired training showed slower acquisition of conditioned food-cup responses in paired training (phase 2), but no retardation in the acquisition of paired learning was observed in Gi+CNO rats with prior unpaired training (*). There was no difference in the acquisition rate of paired learning between Gi-VEH rats without unpaired training and Gi-CNO rats with unpaired training. Error bars represent SEM.

Figure 2F shows food-cup responses to light in the three groups that underwent unpaired training. A two-way repeated-measures ANOVA with group and day as the main factors revealed a significant effect of day (F_(5,125)_ = 7.32, *p* < 0.001), with no significant group effects (F_(2,25)_ = 0.62, *p* = 0.54) or group × day interaction effects (F_(10,125)_ = 1.49, *p* = 0.15), indicating that the animals in the three groups showed decreased food-cup responses during unpaired training.

Figure 2F shows food-cup responses to light during paired training in animals of the four groups, with or without prior unpaired training. A two-way repeated-measures ANOVA with group and day as the main factors revealed a significant effect of day (F_(5,180)_ = 117.27, *p* < 0.001) and group (F_(3,36)_ = 3.95, *p* < 0.05), but no group × day interaction effects (F_(15,180)_ = 1.36, *p* = 0.17). *Post hoc* analyses revealed retardation of acquisition in Gi+VEH and EGFP+CNO groups with prior unpaired training, but not in Gi+CNO rats with prior unpaired training (Figure 2F). These results indicate that chemogenetic LHb inhibition throughout unpaired training eliminates the retardation of acquisition of subsequent excitatory learning following inhibitory learning. Notably, no difference was found between Gi+VEH rats without prior unpaired training and Gi-CNO rats with prior unpaired learning. Interestingly, chemogenetic LHb inhibition throughout paired learning did not prevent retardation in the acquisition of excitatory learning following inhibitory learning (Figure 2-Supplement figure 1).

### A CS that is explicitly unpaired with food passes the summation test of a conditioned inhibitor

The present study also used a summation task to examine a conditioned inhibitor of a CS that was unpaired with food for 6 days. The present conditioning procedure (Figure 3A) was adapted from a conditioned suppression experiment that investigated the effects of explicitly unpaired training of CS and US on the subsequent excitatory conditioning of a compound CS (Rescorla, 1971). Animals in the unpaired-compound paired group received explicitly unpaired training of light and food for 6 days (phase 1) and paired training of compound light-tone with food for the subsequent 6 days (phase 2). Food-cup responses to either light- or tone-alone presentations were measured for the next 2 days (test phase). Two comparison groups were added to determine whether an explicitly unpaired CS with food exhibited inhibitory properties. Rats in the compound paired-alone group received a paired presentation of compound light-tone and food alone. Rats in the paired-compound paired group received light-food pairings in phase 1, followed by compound light-tone and food pairings in phase 2.

**Figure 3.**
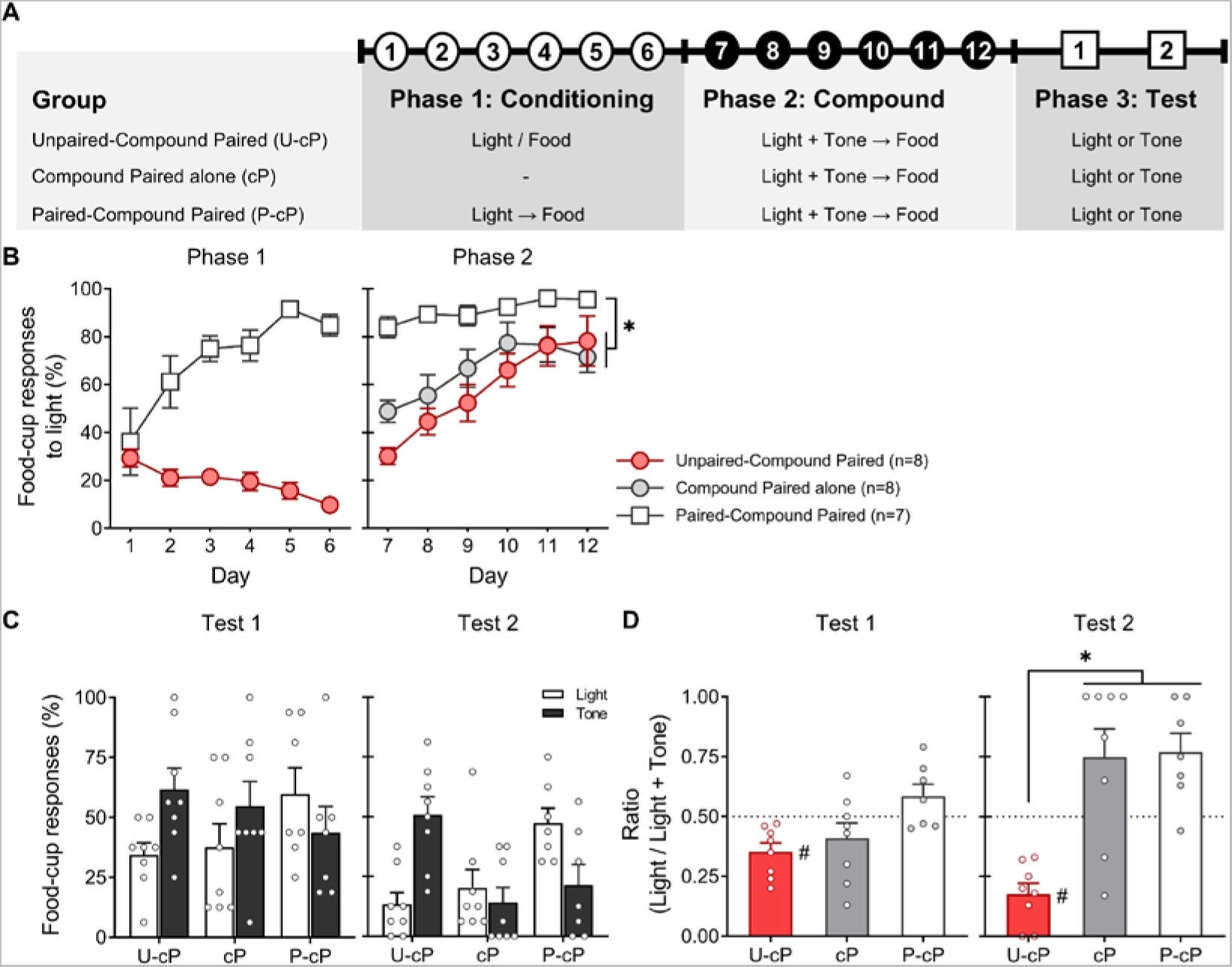
Explicitly unpaired CS with food passes the summation test of inhibition. **(A)** Conditioning procedures. **(B)** In phase 1, rats initially received paired or explicitly unpaired training (light/food). In phase 2, all rats in the two groups underwent compound light-tone and food pairings. Another cohort of rats received compound training alone. The paired-compound paired group exhibited higher food-cup responses to a compound CS than the other groups (*). **(C)** Food-cup responses to either repeated light- or tone-alone presentations, measured in the test phase. **(D)** Inhibitory strength of explicitly unpaired conditioned light, expressed as the light/(light+tone) ratio. (#, denotes significant inhibitory properties of light; one-sample t-test). In test 2, the ratio of light was significantly lower in the unpaired-compound paired group than in the other groups (*). Error bars represent SEM.

Figure 3B shows conditioned food-cup responses in phases 1 and 2. A two-way repeated-measures ANOVA of phase 1 with day and group as the main factors revealed significant effects of day (F_(5,65)_ = 5.47, *p* < 0.001) and group (F_(1,13)_ = 5.91, *p* < 0.001). Moreover, a significant effect of day × training (F_(5,65)_ = 17.19, *p* < 0.001) was noted, indicating a gradual decrease in food-cup responses to light in the unpaired-compound paired group and an increase in food-cup responses to light in the paired-compound paired group. A two-way repeated-measures ANOVA of phase 2 with day and group as the main factors revealed a significant effect of day (F_(5,100)_ = 23.10, *p* < 0.001) and group (F_(1,20)_ = 9.89, *p* < 0.01), and significant day × group interaction effects (F_(2,100)_ = 5.51, *p* < 0.01). *Post hoc* analyses revealed that the paired-compound paired group exhibited higher food-cup responses to a compound CS than the unpaired-compound paired and compound paired alone groups; however, no difference between unpaired-compound paired and compound paired alone groups was observed.

Figure 3 (panels C and D) shows food-cup responses to light or tone and light/(light+tone) ratios in the test phase (see Methods for details). A one-sample t-test of the ratio of each group was performed to determine whether light exerted conditional inhibitory properties, where a value of 0.5 indicates no conditioned inhibitory properties of light. The ratios of tests 1 and 2 in the unpaired-compound paired group were 0.35 and 0.18, respectively, values that were significantly lower than 0.5 (test 1: *t*_(7)_ = -3.98, *p* < 0.01; test 2: *t*_(7)_ = -7.15, *p* < 0.001). These results indicate that light possesses inhibitory properties and demonstrate the occurrence of “superconditioning” of the tone (Rescorla, 1971).

The ratios of tests 1 and 2 in the compound paired-alone group were 0.40 and 0.74, respectively (test 1: *t*_(7)_ = -1.43, *p* = 0.20; test 2: *t*_(7)_ = 2.10, *p* = 0.07). In the paired-compound paired group, the ratios of tests 1 and 2 were 0.58 and 0.77, respectively, the latter of which (test 2) was significantly higher than 0.5 (test 1: *t*_(6)_ = 1.66, *p* = 0.15; test 2: *t*_(6)_ = 3.40, *p* < 0.05), indicating the occurrence of a Kamin blocking effect of light (Kamin, 1968). A one-way ANOVA of inhibition ratio revealed a significant effect of group (test 1: F_(2,20)_ = 5.26, *p* < 0.01; test 2: F_(2,20)_ = 15.25, *p* < 0.001). *Post hoc* analyses revealed no difference between the unpaired-compound paired group and the compound paired alone group in test 1, but did reveal significant differences between the two in test 2.

### Gi-DREADD–mediated LHb inactivation throughout explicitly unpaired training attenuates the inhibitory properties of explicitly unpaired CS, as assessed by the summation test

The behavioral experiment above demonstrated that explicitly unpaired light with food had inhibitory properties. Accordingly, the effects of chemogenetic LHb inhibition on the establishment of an explicitly unpaired conditioned inhibitor were examined (Figure 4A). Specifically, we designed three groups of animals to investigate the effects of reduced LHb activity on the acquisition of an explicitly unpaired conditioned inhibitor: two groups administered hSyn-hM4Di-mCherry-AAV—one provided CNO (Gi+CNO) and one given vehicle (Gi+VEH)—and one control group (EGFP+CNO). Rats received VEH or CNO, as indicated, 30 minutes before explicitly unpaired training (phase 1). No treatments were given during the compound paired training (phase 2) or test phase. Successful infection of the LHb area was confirmed based on the localized expression of mCherry (58.84% ± 3.40%) and EGFP (78.61% ± 43.91%) (Figure 4B–D).

**Figure 4.**
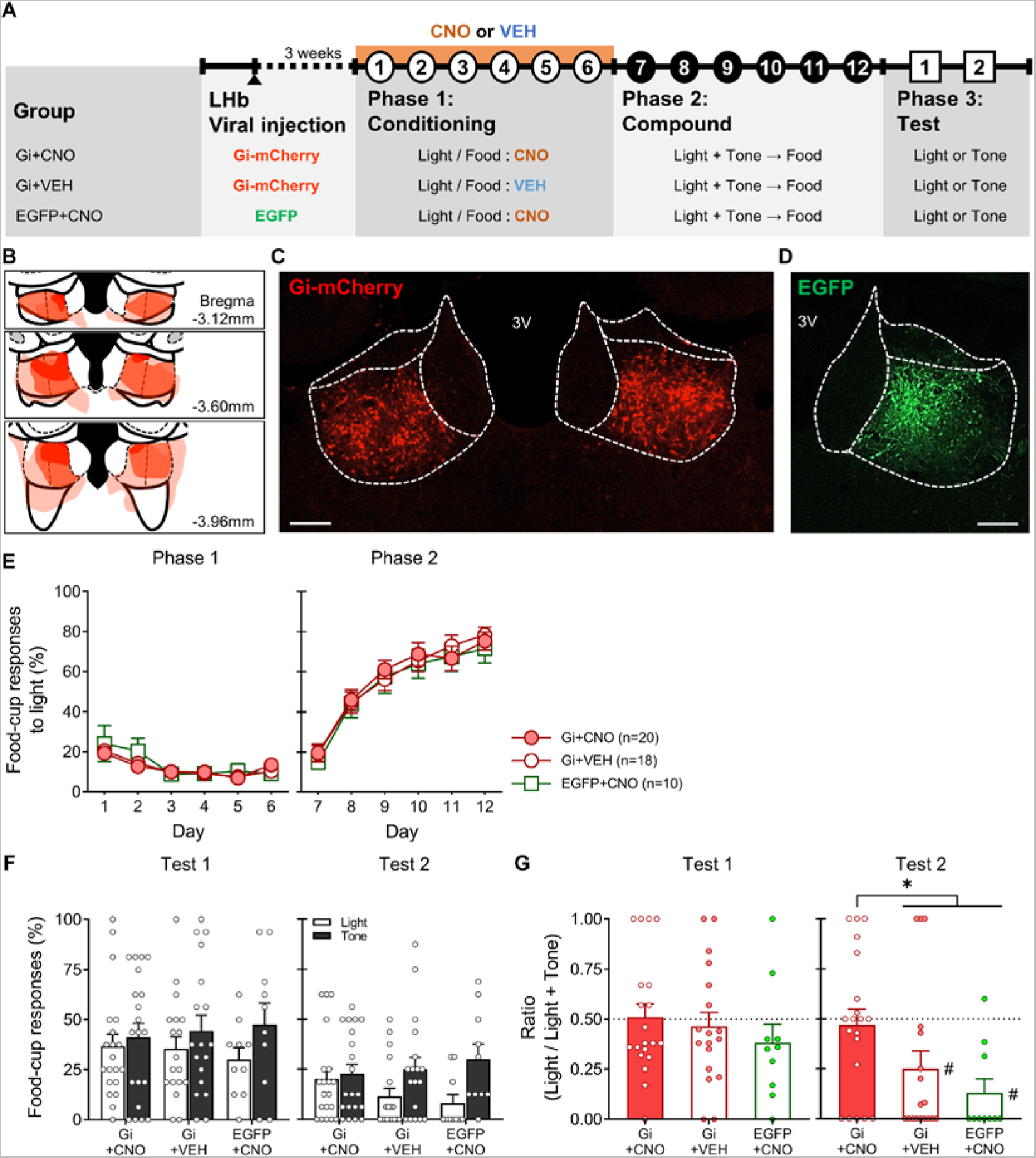
Gi-DREADD–mediated inhibition of the LHb throughout explicitly unpaired training attenuates the inhibitory strength of an explicitly unpaired conditioned stimulus (CS). **(A)** Experimental schedule. **(B)** Histological reconstruction showing the largest (bright), average (medium), and smallest infected (dark) areas. **(C, D)** Representative photomicrographs of the LHb expressing virally transduced hM4Di mCherry or EGFP. Scale bar, 200 µm. 3 V, third ventricle. **(E)** In phase 1, rats initially received explicitly unpaired light and food training. In phase 2, all rats in the three groups underwent compound light-tone and food paired training. **(F)** Food-cup responses to either repeated light- or tone-alone presentations, measured in the test phase. **(G)** The inhibitory strength of explicitly unpaired conditioned light, expressed as the light/(light+tone) ratio. (#, denotes significant inhibitory properties of light; one-sample t-test). In test 2, the inhibitory strength of the Gi-CNO group was attenuated compared with that of Gi+VEH and EGFP+CNO groups (*). Error bars represent SEM.

A two-way repeated-measures ANOVA of phases 1 and 2 (Figure 4E) with day and group as the main factors revealed a significant effect of day (phase 1: F_(5,225)_ = 13.43, *p* < 0.001; phase 2: F_(5,225)_ = 102.17, *p* < 0.001) with no group effect (phase 1: F_(2,45)_ = 0.16, *p* = 0.85; phase 2: F_(2,45)_ = 0.11, *p* = 0.90) or day × group interaction effects (phase 1: F_(10,225)_ = 0.84, *p* = 0.59; phase 2: F_(5,225)_ = 0.48, *p* = 0.90).

Figure 4 (panels F and G) shows food-cup responses to either light or tone and inhibition ratios. The inhibition ratios of tests 1 and 2 in the Gi+VEH group were 0.46 and 0.25, respectively. A one-sample t-test revealed significantly conditional inhibitory properties of light in test 2 (test 1: *t*_(17)_ =-0.52, *p* = 0.61; test 2: *t*_(17)_ = -2.84, *p* < 0.01). The inhibition ratios of tests 1 and 2 in the EGFP+CNO group were 0.38 and 0.13, respectively. A one-sample t-test revealed significant conditional inhibitory properties of light in test 2 (test 1: *t*_(9)_ = -1.28, *p* = 0.23; test 2: *t*_(9)_ = -5.31, *p* < 0.001). However, the inhibition ratios of tests 1 and 2 in the Gi+CNO group were 0.51 and 0.47, respectively, which were not significant in a one-sample t-test (test 1: *t*_(19)_ = 0.15, *p* = 0.88; test 2: *t*_(19)_ = -0.37, *p* = 0.71). A one-way ANOVA of the inhibition ratio revealed no significant effect of group in test 1 (F_(2,45)_ = 0.63, *p* = 0.54), but did reveal a significant group effect in test 2 (F_(2,45)_ = 3.94, *p* = 0.03). *Post hoc* analyses of test 2 revealed that the inhibitory light strength of Gi+VEH and EGFP+CNO groups was significantly greater than that of the Gi+CNO group (*p* < 0.05).

### Chemogenetic LHb excitation facilitates extinction

Repeated presentations of a CS without a US following CS-US learning generates conditioned inhibition of the CS (Pavlov, 1927; Rescorla, 1969a; Calton et al., 1996). Our recent study reported that c-Fos levels in the LHb were increased in rats with extinction training and that extinction was delayed in rats with a neurotoxin-lesioned LHb (Kim et al., 2021). On the basis of these findings, we examined the effect of chemogenetic LHb activation on CR extinction. LHb activity was enhanced by expressing the CNO-activated, experimentally evolved Gq-coupled human M3-muscarinic receptor (hM3Dq)-DREADD, delivered into the LHb via an AAV vector. Four groups—Gq+CNO:Paired-Paired, Gq+CNO:Paired-CS alone, Gq+VEH:Paired-CS alone and EGFP+CNO:Paired-CS alone—were designed to examine the effects of enhanced LHb activation on the extinction of CR and to control for the effects of CNO or transfected gene (see Methods section for details; Figure 5A). Successful infection of the LHb area was confirmed based on the localized expression of mCherry (76.87% ± 3.92%) and EGFP (62.95% ± 8.92%), as shown in Figure 5B. Neuronal LHb activation through intraperitoneal CNO injection was also confirmed by assessing the number of c-Fos–positive signals in rats virally transduced with hM3Dq or EGFP (Figure 5C, D; Group, F_(3,33)_ = 30.41, *p* < 0.001).

**Figure 5.**
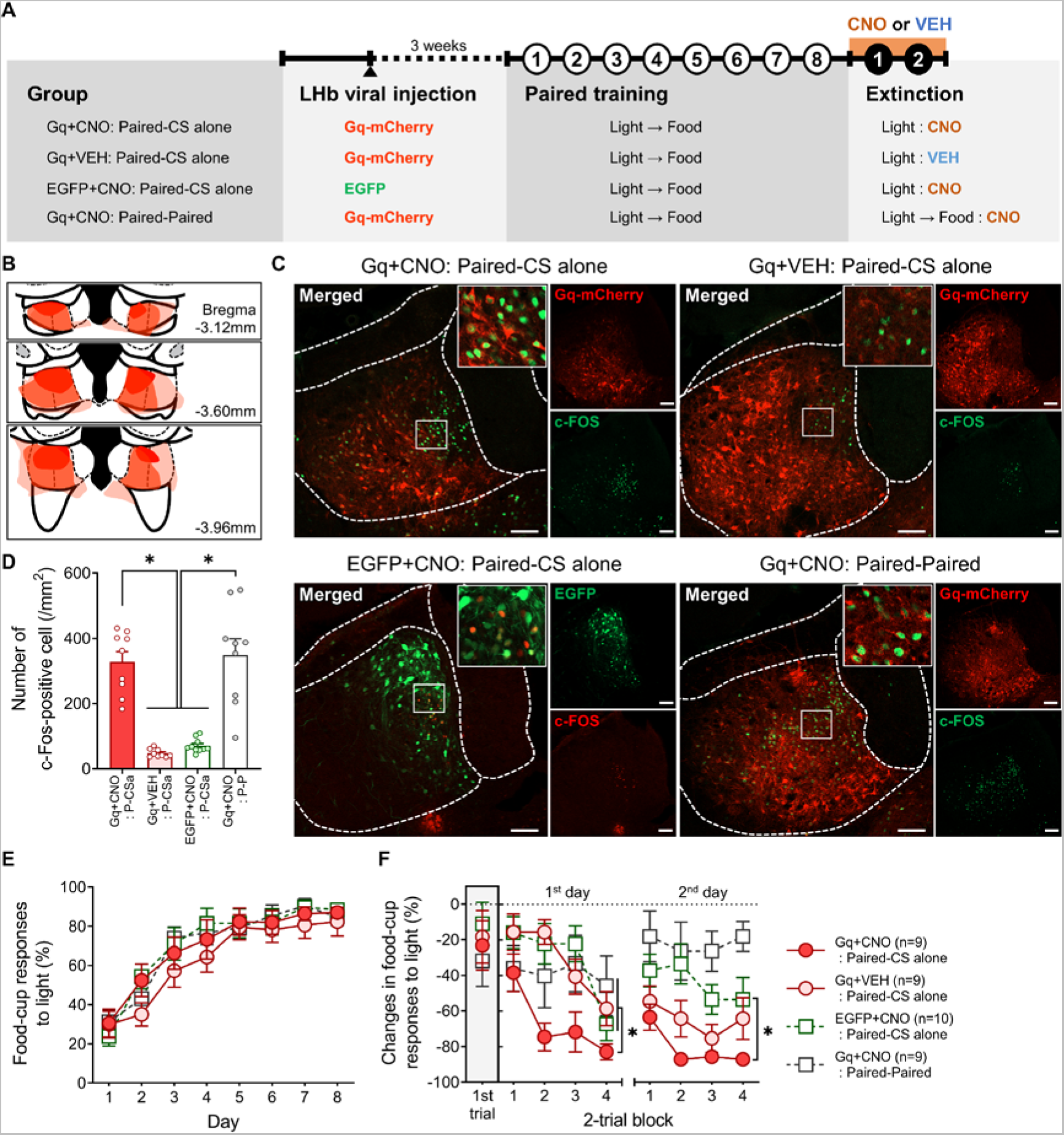
Gq-DREADD–mediated excitation of the LHb facilitates extinction. **(A)** Experimental schedule. **(B)** Histological reconstruction showing the largest (bright), average (medium), and smallest infected areas of the LHb of rats expressing virally transduced hM3Dq mCherry or EGFP. **(C)** Images in the first-row show mCherry-expressing neurons (red) and c-Fos (green) in the LHb of a rat administered CNO or vehicle. Images in the second-row show EGFP-expressing neurons (green) and c-Fos (red) in the LHb of a rat administered CNO. Images at top right and right are magnifications of images in solid-line boxes. Scale bar, 100 µm. **(D)** CNO administration significantly increased c-Fos signals (*). **(E)** Rats in all four groups received eight daily 64-minute light-food paired training trials. All rats in each of the four groups showed comparable acquisition performances. **(F)** Rats in the Gq+CNO:Paired-paired group received additional twice-daily paired training, whereas rats in other groups received twice-daily 64-minute extinction training consisting of eight light-alone presentations. Rats in the Gq+CNO:Paired-CS alone group exhibited a significantly faster decline in conditioned food-cup responses than those in the other groups on the first day (*) and showed a significant decline compared with rats in the EGFP+CNO:Paired-CS alone group on the second day (*#*). In the first trial, performance was equivalent among groups. Error bars represent SEM.

In the acquisition phase, rats in the four groups received daily 64-minute paired training for 8 days (eight trials per day). A two-way repeated-measures ANOVA with day and group as the main factors revealed a significant effect of day (F_(7,231)_ = 90.74, *p* < 0.001), with no group effects (F_(3,33)_ = 0.54, *p* = 0.66) or day × group interaction effects (F_(21,231)_ = 0.94, *p* = 0.54), indicating comparable acquisition of conditioned food-cup responses throughout paired training in all groups (Figure 5E).

In the extinction phase, rats were presented light-alone for 2 days (eight trials per day). Because individual learning curves differed in the acquisition phase, the extinction rate per rat was defined as the change from the averaged conditioned food-cup responses on day 8. A two-way repeated-measures ANOVA of day 1 with a two-trial block and group as the main factors revealed significant block (F_(3,99)_ = 17.07, *p* < 0.001), group (F_(3,33)_ = 3.12, *p* < 0.05), and block × group interaction effects (F_(9,99)_ = 2.68, *p* < 0.01). *Post hoc* analyses showed that rats in the Gq+CNO:Paired-CS alone group exhibited faster declines than the corresponding control groups on the first extinction day (Figure 5F). A two-way repeated-measures ANOVA of day 2 with a two-trial block and group as the main factors revealed significant block (F_(3,99)_ = 3.94, *p* < 0.05) and group effects (F_(3,33)_ = 10.54, *p* < 0.001), but no significant block × group interaction effects (F_(9,99)_ = 0.90, *p* = 0.53). *Post hoc* analyses revealed a significant difference between Gq+CNO:Paired-CS alone and EGFP+CNO:Paired-CS alone groups (*p* < 0.01). However, there was no difference between Gq+CNO:Paired-CS alone and Gq+VEH:Paired-CS alone groups (*p* = 0.08, one-tailed), which might be attributable to a floor effect in the Gq+CNO:Paired-CS alone group.

## Discussion

A series of studies using the two-set strategy (retardation-of-acquisition and summation task) of Rescorla (1969) for diagnosing conditioned inhibition (Rescorla, 1969a) and extinction (Pavlov, 1927; Calton et al., 1996) have demonstrated that the LHb is a crucial part of the brain for acquiring the association of a CS with the nonoccurrence of a US. Retardation of acquisition was abolished in rats with neurotoxic LHb lesions or chemogenetic LHb inactivation. Chemogenetic LHb inhibition also attenuated inhibitory strengths of explicitly unpaired CS, measured with the summation task. Furthermore, chemogenetic LHb activation facilitated the extinction of conditioned food-cup responses. These findings support the interpretation that the LHb mediates the association of a CS with the absence of an appetitive US.

Explicitly unpaired training endows a neutral stimulus (CS) with conditioned inhibitory properties (Rescorla, 1969b; Matzel et al., 1988; Williams and Overmier, 1988; Droungas and LoLordo, 1995). In this study, the inhibitory properties of explicitly unpaired CS light and US food using retardation-of-acquisition and summation tasks were assessed as recommended by Rescorla (Rescorla, 1969a). In the retardation-of-acquisition task, the association of light with the absence of food acquired throughout unpaired training retarded the excitatory association of the same light with food over subsequent paired training. In the summation task, explicitly unpaired training was implemented, and then compound stimuli consisting of an explicitly unpaired CS (light) and a newly added CS (tone) were paired with food. Food-cup responses to light- or tone-alone presentations were measured for the next 2 days. The differences in conditioned food-cup responses between the two CS were expressed as the light/(light+tone) ratio. Significant differences in the ratio between unpaired-compound paired and compound paired alone groups were detected on the second test day but not on the first test day. Repeated CS-alone presentations in the test represent a process called extinction. Therefore, the psychological function underlying extinction might affect the inhibitory properties of the explicitly unpaired CS. For example, a noticeable change in a contingency in the test phase of the summation test would increase attention during the initial trials of extinction (Pearce and Hall, 1980) and thereby reduce the difference between unpaired-compound paired and compound paired alone groups on the first test day. On the other hand, an inhibitory CS generated by repeated CS-alone presentations would add to the inhibitory CS strength established by the prior explicitly unpaired training, leading to greater inhibitory strength of the CS.

The expression levels of c-Fos in the LHb and SNc/VTA of unpaired, paired, and control animals have been reported (Choi et al., 2020), with unpaired animals exhibiting an increase in c-Fos levels in the LHb but a decrease in the SNc/VTA. Interestingly, the opposite pattern of c-Fos expression is observed in the LHb and SNc/VTA of paired animals. Although these observations suggest that the LHb engages in a process that associates a CS with the nonoccurrence of a US, they do not imply causality. An additional study that examined the effects of LHb neurotoxic lesions on the process of associating a CS with the presence/absence of a US reported no lesion effects (Choi et al., 2020). Therefore, in this study, we performed experiments employing retardation-of-acquisition and summation tasks to define the role of the LHb in associating a CS with the nonoccurrence of US (food). First, we examined the effects of LHb lesions on retardation of acquisition. Sham-operated animals showed retardation in the acquisition of conditioned food-cup responses in the paired training following unpaired training. This retardation was not observed in LHb-lesioned animals. Next, we inhibited LHb neuronal activity using Gi-DREADD throughout unpaired learning and evaluated its effects in retardation-of-acquisition and summation tasks. Gi-DREADD–mediated inhibition of LHb neuronal activity prevented retardation of acquisition and attenuated the inhibitory properties of a CS, as assessed by the summation test.

Our recent study observed increased c-Fos levels in the LHb of rats that repeatedly received a CS-alone presentation following CS-US paired training (Kim et al., 2021). In addition, LHb lesions slowed appetitive extinction (Donaire et al., 2019; Kim et al., 2021). Therefore, in contrast to earlier experiments employing an approach based on a loss or reduction in LHb neuronal activity, subsequent experiments designed to evaluate LHb activation effects by Gq-DREADD on the extinction of a CS showed that LHb activation accelerated the decline in conditioned food-cup responses. Notably, conditioned food-cup responses on the first day of the extinction phase decreased in Gq+CNO rats that received paired training instead of CS-alone presentations. These results, taken together with a previous report that optogenetic activation of the LHb-rostromedial tegmental nucleus increased immobility (Proulx et al., 2018), indicate that chemogenetic LHb activation might induce behavioral inhibition. However, all groups, including the Gq+CNO group, showed comparable food-cup responses during the pre-CS period, and there were no between-group differences. Therefore, these results indicate that the accelerated declines in conditioned food-cup responses in Gq+CNO rats are not attributable to behavioral inhibition.

Pavlov (1927) initially proposed a neural mechanism of conditioned inhibition in which specific regions of the cerebral cortex mediate conditioned excitation and inhibition in a mutually exclusive, antagonistic manner. Numerous neurobehavioral reports employing a loose definition of conditioned inhibition have studied neural substrates of conditioned inhibition (Sosa and Ramirez, 2019; Sosa et al., 2021). In further support of Pavlov’s theory, it has been reported that dopamine neurons in the midbrain (not the originally predicted cortical region) are activated by reward-predicting stimuli while being inhibited by reward omission-predicting stimuli (Schultz, 1998; Tobler et al., 2003; Chang et al., 2018). In contrast, the models proposed by Konorski and Pearce-Hall posited separate mechanisms for conditioned excitation and inhibition (Konorski, 1967; Pearce and Hall, 1980). Reports that neuronal responses in midbrain dopamine and LHb neurons reflect reward prediction errors (Matsumoto and Hikosaka, 2007), together with the present findings and the results of our previous studies on c-Fos (Choi et al., 2020) and LHb lesions (Laurent et al., 2017; Kim et al., 2021), suggest that the LHb, in tandem with the regulation of dopamine neurons, mediates conditioned inhibition. This idea is consistent with the models of Konorski and Pearce-Hall.

Optimal guidance for approach and avoidance behaviors for an organism may depend on the normal complementary functions of the LHb and midbrain dopaminergic systems for reward and anti-reward information (Trusel et al., 2019). Indeed, contingency learning in complex environments can be used to optimize adaptive behavior and flexible responses under changing conditions (Stopper et al., 2014; Mizumori and Baker, 2017). In this context, recent attention has focused on the maladaptive function of the LHb in psychiatric depression, a view supported by imaging and electrophysiological studies reporting that the LHb is hyperactivated in depressed patients and animal models of depression (Morris et al., 1999; Shumake et al., 2003; Li et al., 2011; Park et al., 2017). The active interest in the LHb highlights its role in the broad integration of forebrain anatomical inputs and the control of critical ascending systems, such as the dopaminergic system, in the complex functions of decision-making and choice behavior. The current study presents evidence for a role for the LHb in conditioned inhibition—a fundamental process in associative learning.

## Materials and Methods

### Animals

The animals were procured from Taconic (Seoul, South Korea) and included 290 male naïve 12 weeks-old male Sprague–Dawley rats (specific-pathogen-free). They were housed individually in a climate-controlled vivarium under a 12:12 h light-dark cycle (lights on from 08:00 h to 20:00 h) at a controlled temperature of (22 ± 1)°C and humidity of (50 ± 10)%. Water and food were provided ad libitum throughout the experiments, but the latter was restricted during the food restriction period of behavioral experiments. All animal experiments were approved and performed in accordance with the guidelines of the Institutional Animal Care and Use Committee of the Konkuk University (KU18019, KU19096, KU20070, KU20137, KU20138).

### Apparatus

The behavioral training apparatus, measuring 22.9 × 20.3 × 28 cm, consisted of four chambers (Coulbourn Instruments, Holliston, MA, USA). Each chamber had front and back walls, a top made of aluminum, transparent acrylic sides, and a floor made of stainless-steel rods (0.48 cm in diameter spaced 1.9 cm apart). A food cup fitted with a photocell sensor for detecting head entries was placed 2 cm above the floor at the front wall center. A jeweled 2-W lamp, mounted on the chamber’s front panel, 22 cm above the food cup, served as the visual cue source. A speaker mounted on the chamber’s back wall on the opposite side from the visual cue generated a tone (70 dB). Each chamber was enclosed in a sound-attenuated box, where a 10-W red light provided constant dim illumination, and ventilation fans provided masking noise (65 dB). A digital video camera was mounted within each box, and the behavior of the subjects was recorded during behavioral training.

### Behavioral observation procedure

All behavioral training procedures were adapted and modified from our previous study(Choi et al., 2020). All rats were gradually reduced to 85% of their free-feeding body weights and maintained at that weight by providing a premeasured amount of rat chow each day in the late afternoon. Food restriction was started a week before the behavioral experiments and maintained until the end of the experiments. The shaping was started in 12-week-old rats. Sixteen deliveries of two 45-mg food pellets (Bio-serv, Flemington, New Jersey, USA) were given on a variable-time of 4-min schedule ranging from 2 to 6 min in a single day to acclimatize rats to food delivery. Next, 16 10-s light-alone trials, with intertrial intervals of an average of 4 min, were presented before the conditioned stimulus (CS)-unconditioned stimulus (US) pairings or CS-US unpairing for the habituation of the orienting rearing to light. Additional same habituation procedure for tone was conducted on another day for the summation task.

### Conditioned response analysis

Behavioral data were automatically recorded in the log data file using the Graphic State 4 program (Coulbourn Instruments). The primary measure of conditioning was that the photocell in the recessed food cup was broken (indicating the presence of the rat’s head in those cups) at 1.25-s intervals during a CS presentation. Because these behaviors occur predominantly a few seconds immediately before delivery of the reinforcer(Holland, 1977; Holland, 1984), the behaviors that occurred during the last 5 s of the 10-s CS presentation were analyzed (Gallagher et al., 1990; Choi et al., 2020). We also measured the food-cup responses in the 5-s interval before a CS presentation (pre-CS). In the extinction task, changes in the food-cup responses were expressed as a decline from the food-cup responses in the last acquisition day per animal.

### Retardation-of-acquisition and summation task procedure

Unpaired training was continued for 6 days in which the same number of lights (CS) and food pellets (US) were pseudorandomly presented using an explicitly unpaired procedure such that they were never exhibited at the same time, with the provision that an identical stimulus did not appear for three consecutive times (0% CS-US contiguity and negative contingency). The paired training was performed for 6 days in which eight 10-sec presentations of the light stimulus were immediately followed by food intake, with variable intertrial intervals with an average of 8 min (100% CS-US contingency). These paired procedures supported the development of associatively food-cup behavior that was not observed with unpaired CS-US presentations(Gallagher et al., 1990; Han et al., 1997).

In the retardation-of-acquisition task, rats received 64-min unpaired light-food training for 6 days (phase 1), followed by 64-min paired light-food training for the next 6 days (phase 2). Another cohort of rats received 64-min paired training alone for 6 days.

In the summation task, the rats received 64-min unpaired light-food training for 6 days (phase 1) and subsequent for 6 days 64 min compound (light+tone)-paired training for 6 days (phase 2). A cohort of rats received 64-min paired light-food training for 6 days (phase 1), followed by 64-min compound-paired training for 6 days (phase 2). Furthermore, the rats in the comparison group received 64-min compound-paired training alone for 6 days. In the test phase of the task, the food-cup responses to either repeated light- or tone-alone presentations were measured during 2 days in which the four 10-sec lights and tones were pseudorandomly presented with variable intertrial intervals and an average of 8 min such that an identical stimulus did not appear for three consecutive times.

### Extinction task procedure

The rats assigned to the extinction experiment received 64-min paired training for 8 days, as described above. Next, 64-min extinction with eight light-alone presentations (variable intertrial intervals with an average of 8 min) or an additional 64-min paired training was followed for 2 days.

### Lesions of the lateral habenula (LHb) and histology

The rats were mounted on a stereotaxic frame (Kopf Instruments, Tujunga, CA, USA) fitted with an isoflurane gas anesthesia system. The scalp was incised and retracted, and small burr holes were drilled on the skull above the LHb. The rats received bilateral lesions in the LHb with ibotenic acid (5 µg/µl in 0.01 M PBS, Abcam, Cambridge, UK). The coordinates for the LHb (AP −3.3 mm, ML ±0.7 mm, DV −5.1 mm) were modified from a previous study(Gifuni et al., 2012). Ibotenic acid or an appropriate vehicle at a volume of 0.3 µl/site was delivered in 6 min using a Hamilton microsyringe. The injection cannula was left in place for 6 min and then removed. Every sixth section with brain regions, including the LHb, was mounted on the slides for Nissle-stain to verify the neurotoxic lesions of the LHb. Brain sections from lesioned rats were imaged using the Eclipse Ni uplight microscope (Nikon, Tokyo, Japan) and Progres® camera (Jenoptik, Jena, Germany) interfaced to a computer. Analyses of the lesioned areas were performed using NIH ImageJ.

### Chemogenetic modulation

The rats received viral injections into the LHb using the same stereotaxic surgery method and the coordinates described in the lesion method. Adeno-associated virus (AAV) vectors of inhibitory Gi-coupled designer receptors exclusively activated by designer drugs (DREADDs) (hSyn-hM4Di-mCherry-AAV5 half dilution of stocks, 8.6 × 10^12^ GC/mL titer, or hSyn-hM4Di-mCherry-AAV2 1:10 dilution of stocks, 1.3 × 10^13^ GC/mL titer, plasmid #50475 from Addgene, Watertown, MA, USA), excitatory Gq-coupled DREADDs (hSyn-hM3Dq-mCherry-AAV1/2 viral stocks, 4.0 × 10^11^ GC/mL titer, plasmid #50474 from Addgene), and control construct (hSyn-enhanced green fluorescent protein (EGFP)-AAV2 1:10 dilution of viral stocks, 8.6 × 10^12^ GC/mL titer, or hSyn-EGFP-AAV5 1:20 dilution of stocks, 1.3 × 10^13^ GC/mL titer, plasmid #50465 from Addgene) were injected into the LHb (1.0 μL/site) at the same coordinates as the lesioned coordinates. All injections were administered at a 0.1 μL/min infusion rate using a 28-gauge needle attached to a 2 μL or 5 μL Hamilton microsyringe (Sigma-Aldrich, St. Louis, MO, USA). The injection cannula was left in place for 15 min after the infusions before retracting slowly. The rats were given 3 weeks of postoperative recovery before commencing behavioral experiments.

### Clozapine-N-oxide treatment

Clozapine-*N*-oxide (CNO; 1.0 mg/kg) (Tocris, Bristol, UK), dissolved in 0.25% dimethyl sulfoxide or 0.9% sterile saline vehicle solution, was administered intraperitoneally to rats 30 min before unpaired training or the extinction procedure. The rats for the c-Fos experiment were perfused 30 min after the last light presentation or the last paired trial.

### Immunohistochemistry

Upon completion of the behavioral testing, rats were deeply anesthetized with isoflurane and transcardially perfused with cold phosphate-buffered saline (PBS) followed by 4% paraformaldehyde in PBS to determine the locus and extent of the DREADD expression. The brains were removed, post-fixed in the same paraformaldehyde solution for 3 days, and submerged in a 30% sucrose solution for 4 days. A microtome was used to cut the brains into 30–40 μm coronal sections. The brain sections of the rats with Gi-DREADDs were incubated first in a blocking solution (10% fetal horse serum and 0.15% Triton X-100 in PBS) for 1 h at 24 ± 1°C and then with primary rabbit anti-mCherry antibody (1:1000, Abcam, ab167453, Cambridge, UK) overnight at room temperature. After washing in PBS containing 0.15% Triton X-100 (PBS-T), the sections were incubated for 1 h at room temperature with Alexa Fluor 488-conjugated donkey anti-rabbit secondary antibody (1:200; Invitrogen, A-21206, Waltham, MA, USA). The Gi-DREADD or stained brain sections were mounted on resin-coated slides and dried slightly. Finally, the sections were coverslip-mounted with ProLong diamond anti-fade reagent (Invitrogen).

The c-Fos expression was measured 2 h after intraperitoneal (IP) injections of vehicle (VEH) or CNO in the rats with Gq-DREADDs to verify that the Gq-DREADDs effectively activated the LHb neurons. The rats with Gq-DREADDs or EGFP control virus received an IP injection of CNO or VEH 30 min before the extinction procedure. Subsequently, they were sacrificed 90 min after commencing the extinction procedure. All perfusion and staining procedures described above were used until incubation with primary rabbit anti-c-Fos antibody (1:500; Santa Cruz, sc-52, Dallas, TX, USA), after which the sections were incubated for 1 h with secondary Alexa Fluor 488 donkey anti-rabbit antibodies (1:200, Invitrogen, A-21206) or Alexa Fluor 568 donkey anti-rabbit antibodies (1:200, Invitrogen, A10042). Sections were mounted on resin-coated slides and dried slightly. Finally, the mounted sections were cover-slipped mounted with a ProLong gold anti-fade reagent (Invitrogen).

Images were obtained using an LSM 900 confocal laser scanning microscope (Carl Zeiss, Oberkochen, Germany) or an Eclipse Ni fluorescence microscope (Nikon). mCherry immunofluorescence was measured to determine the location and area of viral infection. The c-Fos^+^ fluorescent signals in the LHb from −2.9 mm to −4.0 mm (magnified ×100) were analyzed using ImageJ software (NIH). The c-Fos^+^ signals were counted bilaterally in at least six sections (every fourth) per animal.

### Statistical analyses

A two- or three-way repeated and a one-way ANOVA was used to analyze the behavioral data and c-Fos levels, respectively. LSD *post-hoc* tests were performed to examine group differences, if necessary. The food-cup responses to either light or tone in the test phase of the summation task were measured, and then a comparison of the food-cup responses to the two stimuli was expressed as inhibition ratio (light/[light + tone]) per animal. The one-sample t-test (two-tailed significance with the test value setting of 0.5, no conditioned inhibitory properties of light) was employed to analyze the inhibition ratios from days 1 and 2. All data are expressed as the mean ± SEM. Values of *p* < 0.05 were considered significant unless otherwise specified (SPSS statistics 25 programs; IBM, Armonk, NY, USA). Prism 8 software (GraphPad Software, San Diego, CA, USA) was used for preparing graphical figures.

## Acknowledgments

This work was supported by the National Research Foundation of Korea(NRF) grant funded by the Korea government (2020R1A2C1005148 and 2020M3E5D9080734 to J.S.H; NRF-2020R1A6A3A01099505 to D.H.K). This paper was written as part of Konkuk University’s research support program for its faculty on sabbatical leave in 2021.

## Author Contributions

D.H.K. performed Gi-summation, extinction, and data curation. I.B.J. performed LHb lesions in retardation and latent inhibition, Gi-retardation, and data curation. N.H.K. generated behavioral experiments of the summation task. Y.J.J. carried out the surgery of the lesion experiment. B.R.C. contributed to conceptualization and data curation. M.G. wrote and edited the manuscript; J.S.H. contributed to the conceptualization and supervision of the study, fund acquisition, drafting of the original manuscript, editing of the manuscript, and project administration.

## Supplementary Materials List

**Figure 1-Supplement figure 1.**
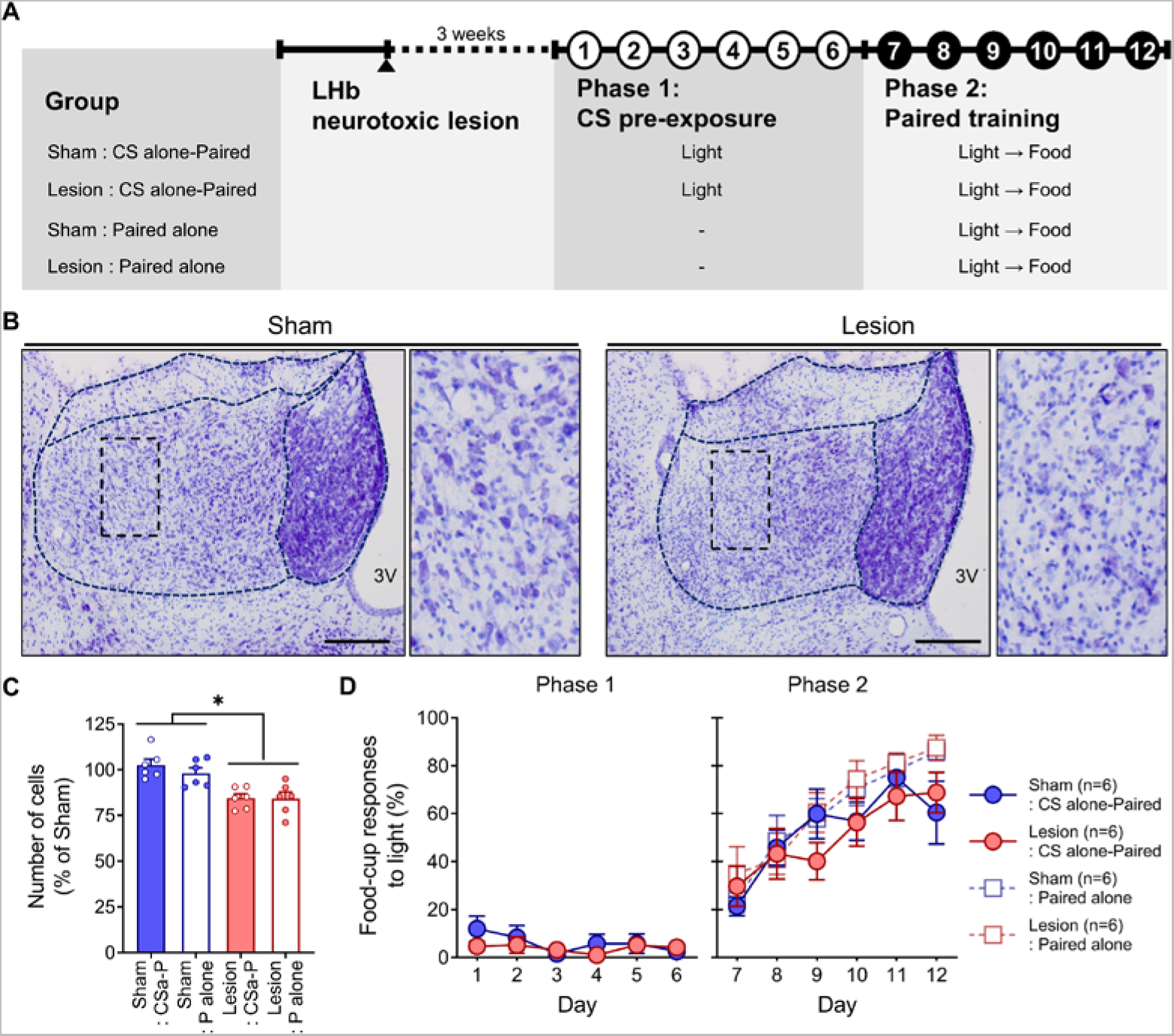
No effects of LHb lesion on latent inhibition. **(A)** Experimental schedule. In each of the six daily 64-minute CS preexposure days, rats in the two groups received eight 10-second light presentations, followed by six daily 64-minute acquisition phases with eight light-food paired presentations. Another cohort of sham-operated and LHb-lesioned rats (sham:paired and lesion:paired) received the same paired training alone without CS preexposure. **(B)** Photomicrographs show Nissl-stained control (left) and neurotoxin-lesioned LHb (right), with evident loss of neurons. Magnified images of the dotted-boxed area are presented at right. Scale bar, 200 µm. 3 V, third ventricle. **(C)** Two-way ANOVA of cell densities with training and lesion showed a significant lesion effect (F_(1,20)_ = 27.83, *p* < 0.001) with no training effect (F_(1,20)_ = 0.64, *p* = 0.43) or lesion × training interaction effects (F_(1,20)_ = 0.53, *p* = 0.48). **(D)** No differences in food-cup responses were found between sham and lesion groups during CS preexposure (phase 1; lesion, F_(1,10)_ = 0.68, *p* = 0.43; day F_(5,50)_ = 1.16, *p* = 0.34; lesion × day interaction, F_(5,50)_ = 0.70, *p* = 0.63). In phase 2, CS preexposure (48 trials) did not significantly retard acquisition of subsequent paired learning (F_(1,20)_ = 3.78, p = 0.07). There was no effect of LHb lesion (F_(1,20)_ = 0.04, *p* = 0.84) or interaction between the two (F_(1,20)_ = 0.53, *p* = 0.47). Error bars represent SEM.

**Figure 2-Supplement figure 1.**
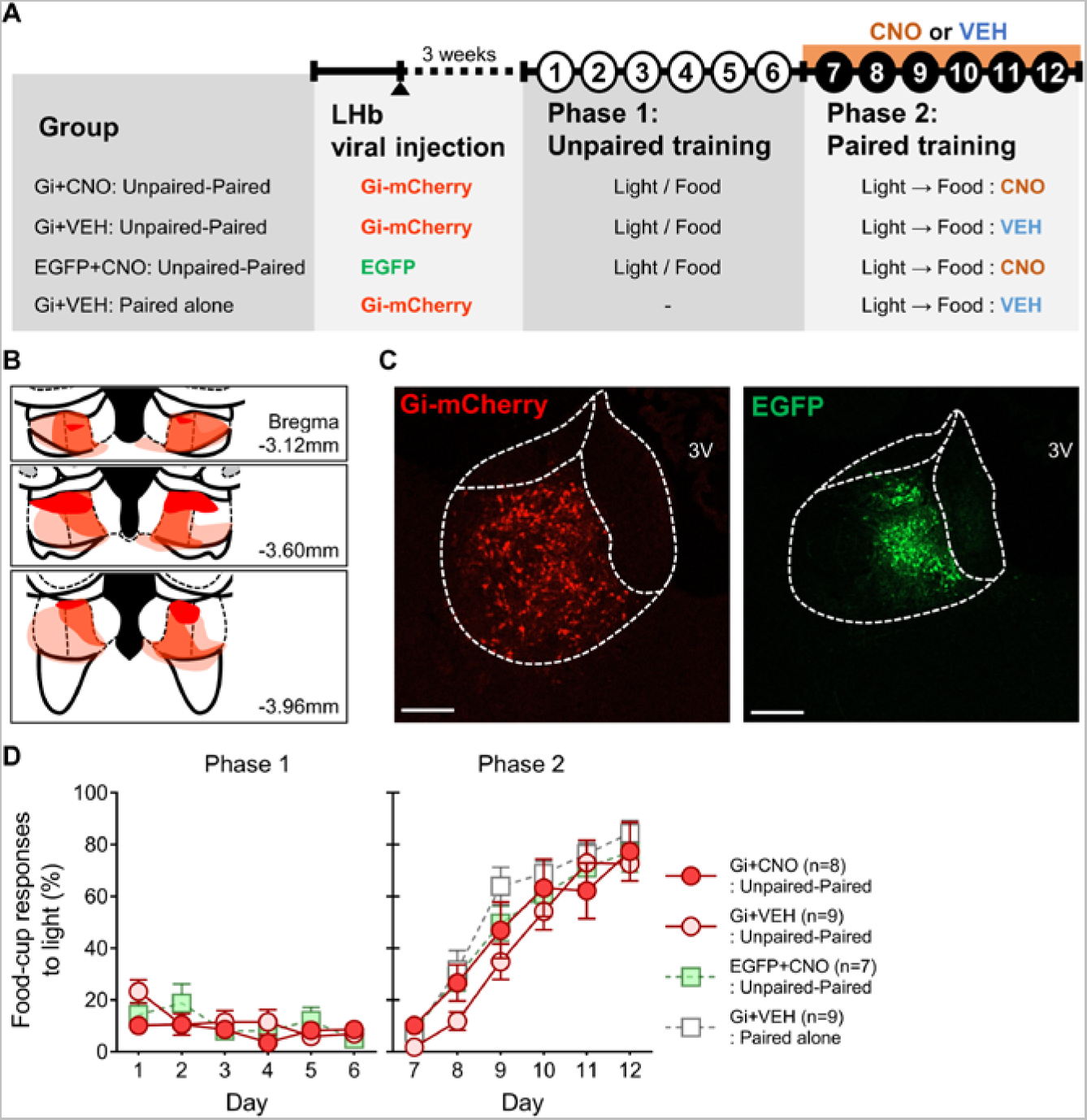
Animals with Gi-DREADD–mediated inactivation of the LHb throughout paired training exhibit retardation of acquisition. **(A)** Experimental schedule. **(B, C)** Histological reconstruction showing the largest (bright), average (medium), and smallest infected (dark) areas and representative photomicrographs of LHb expressing virally transduced hM4Di or EGFP. Scale bar, 200 µm. Successful infection of the LHb area was confirmed based on the localized expression of mCherry (36.15% ± 4.46%) or EGFP (24.80% ± 9.75%). **(D)** Left: Conditioned food-cup responses during unpaired training (phase 1). A two-way repeated-measures analysis of variance (ANOVA) with group and day as the main factors revealed a significant effect of day (F_(5,105)_ = 3.83, *p* < 0.01) with no significant group effects (F_(2,21)_ = 0.56, *p* = 0.58) or interaction group × day effects (F_(10,105)_ = 1.71, *p* = 0.09), indicating that animals in the three groups show decreased food-cup responses during unpaired training. Right: Food-cup responses to light during paired training with or without prior training (phase 2). A two-way repeated-measures ANOVA with group (Gi+VEH rats without prior unpaired training vs. Gi+VEH rats with prior unpaired training) and day as main factors revealed significant effects of day (F_(5,80)_ = 84.28, p < 0.001) and group (F_(1,16)_ = 5.74, p < 0.05), but no group × day interaction effects (F_(5,80)_ = 2.16, p= 0.07). These results indicate that animals in Gi+VEH rats with prior unpaired training show retardation in the acquisition of conditioned food-cup responses throughout paired training. An examination of the effect of chemogenetic LHb inhibition during paired training on retardation of acquisition by two-way repeated-measures ANOVA with group (three groups with prior unpaired training) and day as main factors revealed a significant effect of day (F_(5,105)_ = 66.05, p < 0.001), but no significant group effects (F_(2,21)_ = 0.62, p = 0.55) or group × day interaction effects (F_(10,105)_ = 0.79, p = 0.64). These results indicate that chemogenetic LHb inhibition throughout paired training does not affect the retarded acquisition of subsequent excitatory learning following inhibitory learning. Error bars represent SEM.

**Supplement table 1.** Food-cup behaviors during the 5 secs immediately before CS presentations

**Retardation - LHb lesion (figure 1)**
| Group | Day | Phase 1: Unpaired conditioning |  |  |  |  |  | Phase 2: Paired conditioning |  |  |  |  |  |
| --- | --- | --- | --- | --- | --- | --- | --- | --- | --- | --- | --- | --- | --- |
|  |  | 1 | 2 | 3 | 4 | 5 | 6 | 7 | 8 | 9 | 10 | 11 | 12 |
| Sham<br>: Unpaired-Paired | n=6 | 27.08<br>± 5.27 | 16.67<br>± 5.08 | 15.10<br>± 6.53 | 7.81<br>± 2.65 | 6.77<br>± 2.20 | 6.25<br>± 2.91 | 13.54<br>± 7.16 | 6.25<br>± 2.91 | 7.81<br>± 3.30 | 5.21<br>± 2.38 | 11.98<br>± 4.07 | 3.13<br>± 1.61 |
| Lesion<br>: Unpaired-Paired | n=6 | 25.52<br>± 4.22 | 16.15<br>± 2.60 | 15.10<br>± 5.19 | 5.73<br>± 1.49 | 12.50<br>± 2.42 | 10.94<br>± 2.65 | 15.63<br>± 6.60 | 10.42<br>± 2.64 | 8.33<br>± 3.49 | 9.90<br>± 7.05 | 14.58<br>± 5.74 | 18.23<br>± 3.99 |
| Sham<br>: Paired alone | n=6 | - | - | - | - | - | - | 19.27<br>± 6.67 | 9.38<br>± 2.55 | 18.75<br>± 4.19 | 13.54<br>± 4.32 | 13.54<br>± 5.15 | 12.50<br>± 5.04 |
| Lesion<br>: Paired alone | n=6 | - | - | - | - | - | - | 17.71<br>± 7.77 | 15.10<br>± 3.37 | 10.94<br>± 4.62 | 18.23<br>± 9.42 | 25.52<br>± 6.11 | 13.54<br>± 6.69 |

**Retardation - inhibition of LHb with DREADDs (figure 2)**
| Group | Day | Phase 1: Unpaired conditioning |  |  |  |  |  | Phase 2: Paired conditioning |  |  |  |  |  |
| --- | --- | --- | --- | --- | --- | --- | --- | --- | --- | --- | --- | --- | --- |
|  |  | 1 | 2 | 3 | 4 | 5 | 6 | 7 | 8 | 9 | 10 | 11 | 12 |
| Gi+CNO<br>: Unpaired-Paired | n=10 | 17.81<br>± 2.87 | 17.50<br>± 4.85 | 14.69<br>± 3.09 | 5.63<br>± 1.67 | 7.19<br>± 1.87 | 4.69<br>± 2.25 | 13.13<br>± 3.29 | 13.13<br>± 4.24 | 9.38<br>± 3.39 | 13.13<br>± 3.89 | 10.94<br>± 3.62 | 7.50<br>± 3.17 |
| Gi+VEH<br>: Unpaired-Paired | n=11 | 23.58<br>± 6.34 | 25.28<br>± 7.66 | 21.31<br>± 6.35 | 18.47<br>± 4.81 | 15.06<br>± 4.71 | 14.20<br>± 6.33 | 12.78<br>± 4.07 | 10.51<br>± 4.05 | 16.76<br>± 5.35 | 13.64<br>± 4.16 | 9.66<br>± 3.28 | 19.89<br>± 6.61 |
| EGFP+CNO<br>: Unpaired-Paired | n=8 | 25.00<br>± 5.41 | 14.84<br>± 4.90 | 14.45<br>± 5.01 | 25.78<br>± 7.74 | 8.98<br>± 2.39 | 10.94<br>± 3.07 | 11.72<br>± 4.49 | 9.77<br>± 3.56 | 13.28<br>± 3.63 | 10.16<br>± 3.82 | 14.06<br>± 5.32 | 11.72<br>± 4.45 |
| Gi+VEH<br>: Paired alone | n=11 | - | - | - | - | - | - | 16.48<br>± 4.66 | 17.61<br>± 3.51 | 16.19<br>± 6.50 | 9.09<br>± 3.73 | 12.22<br>± 4.24 | 12.50<br>± 3.93 |

Summation test (figure 3)
| Group | Day | Phase 1: Pavlovian conditioning |  |  |  |  |  | Phase 2: Compound conditioning |  |  |  |  |  | Phase 3: Test |  |  |  |
| --- | --- | --- | --- | --- | --- | --- | --- | --- | --- | --- | --- | --- | --- | --- | --- | --- | --- |
|  |  | 1 | 2 | 3 | 4 | 5 | 6 | 7 | 8 | 9 | 10 | 11 | 12 | 13<br>(Light) | 13<br>(Tone) | 14<br>(Light) | 14<br>(Tone) |
| Unpaired-Compound Paired | n=8 | 30.08<br>± 4.65 | 24.61<br>± 5.36 | 26.56<br>± 8.48 | 25.00<br>± 4.87 | 20.31<br>± 3.96 | 23.05<br>± 2.57 | 30.86<br>± 8.96 | 20.31<br>± 6.25 | 20.70<br>± 4.53 | 19.14<br>± 4.51 | 25.00<br>± 3.64 | 17.97<br>± 2.49 | 8.59<br>± 2.88 | 8.59<br>± 5.14 | 3.91<br>± 2.62 | 18.75<br>± 7.83 |
| Compound Paired alone | n=8 | - | - | - | - | - | - | 21.48<br>± 3.85 | 21.09<br>± 5.21 | 14.06<br>± 2.77 | 15.23<br>± 5.43 | 14.84<br>± 4.37 | 22.27<br>± 6.85 | 16.41<br>± 6.46 | 13.28<br>± 5.84 | 10.16<br>± 4.56 | 4.69<br>± 2.29 |
| Paired-Compound Paired | n=7 | 20.54<br>± 9.74 | 26.79<br>± 7.31 | 24.55<br>± 6.41 | 12.95<br>± 1.86 | 13.39<br>± 5.82 | 17.41<br>± 7.08 | 14.73<br>± 8.51 | 17.41<br>± 9.81 | 24.55<br>± 12.53 | 24.55<br>± 10.68 | 18.75<br>± 9.07 | 17.86<br>± 7.68 | 15.18<br>± 9.13 | 16.96<br>± 7.05 | 7.14<br>± 4.61 | 13.39<br>± 6.88 |

Summation test - inhibition of LHb with DREADDs (figure 4)
| Group | Day | Phase 1: Pavlovian conditioning |  |  |  |  |  | Phase 2: Compound conditioning |  |  |  |  |  | Phase 3: Test |  |  |  |
| --- | --- | --- | --- | --- | --- | --- | --- | --- | --- | --- | --- | --- | --- | --- | --- | --- | --- |
|  |  | 1 | 2 | 3 | 4 | 5 | 6 | 7 | 8 | 9 | 10 | 11 | 12 | 13<br>(Light) | 13<br>(Tone) | 14<br>(Light) | 14<br>(Tone) |
| Gi+CNO | n=20 | 22.81<br>± 4.01 | 18.75<br>± 3.43 | 13.91<br>± 2.99 | 16.88<br>± 2.97 | 12.50<br>± 2.19 | 14.84<br>± 3.28 | 14.06<br>± 2.70 | 9.22<br>± 2.23 | 14.53<br>± 2.55 | 10.63<br>± 2.49 | 9.53<br>± 2.46 | 10.63<br>± 3.23 | 13.75<br>± 3.92 | 8.75<br>± 2.54 | 7.81<br>± 2.79 | 15.94<br>± 4.31 |
| Gi+VEH | n=18 | 23.44<br>± 4.07 | 16.67<br>± 3.15 | 12.15<br>± 2.62 | 11.11<br>± 2.60 | 14.41<br>± 2.62 | 19.10<br>± 3.96 | 10.76<br>± 3.23 | 9.20<br>± 2.27 | 9.72<br>± 1.85 | 13.19<br>± 2.90 | 9.55<br>± 2.15 | 8.68<br>± 2.76 | 11.81<br>± 3.81 | 5.56<br>± 2.20 | 5.90<br>± 2.55 | 8.68<br>± 3.12 |
| EGFP+CNO | n=10 | 30.31<br>± 9.03 | 18.13<br>± 4.08 | 11.25<br>± 4.25 | 12.19<br>± 5.18 | 11.56<br>± 3.81 | 19.06<br>± 4.26 | 14.69<br>± 6.29 | 17.81<br>± 4.19 | 18.13<br>± 5.64 | 14.69<br>± 4.89 | 19.06<br>± 5.54 | 15.00<br>± 7.06 | 10.00<br>± 3.51 | 11.25<br>± 5.50 | 5.00<br>± 3.20 | 6.88<br>± 3.01 |

Extinction - activation of LHb with DREADDs (figure 5)
| Group |  | Day or<br>[Block]<sup>#</sup> | Paired conditioning |  |  |  |  |  |  |  | Extinction |  |  |  |  |  |  |  |
| --- | --- | --- | --- | --- | --- | --- | --- | --- | --- | --- | --- | --- | --- | --- | --- | --- | --- | --- |
|  |  |  |  |  |  |  |  |  |  |  | 1<sup>st</sup> session |  |  |  | 2<sup>nd</sup> session |  |  |  |
|  |  |  | 1 | 2 | 3 | 4 | 5 | 6 | 7 | 8 | [1] | [2] | [3] | [4] | [5] | [6] | [7] | [8] |
| Gq+CNO<br>: Paired-CS alone | n=9 |  | 24.65<br>± 3.10 | 27.78<br>± 4.84 | 26.04<br>± 5.98 | 13.19<br>± 3.45 | 22.57<br>± 3.92 | 17.71<br>± 5.18 | 14.58<br>± 5.00 | 11.46<br>± 2.60 | 22.22<br>± 10.78 | 4.17<br>± 4.17 | 9.72<br>± 5.81 | 0.00<br>± 0.00 | 12.50<br>± 6.91 | 0.00<br>± 0.00 | 0.00<br>± 0.00 | 0.00<br>± 0.00 |
| Gq+VEH<br>: Paired-CS alone | n=9 |  | 25.00<br>± 7.98 | 27.43<br>± 4.06 | 18.75<br>± 4.57 | 14.93<br>± 7.34 | 13.19<br>± 3.92 | 12.15<br>± 3.70 | 8.68<br>± 3.03 | 11.81<br>± 3.07 | 12.50<br>± 5.89 | 5.56<br>± 5.56 | 0.00<br>± 0.00 | 9.72<br>± 5.81 | 5.56<br>± 3.03 | 5.56<br>± 5.56 | 0.00<br>± 0.00 | 11.11<br>± 7.35 |
| EGFP+CNO<br>: Paired-CS alone | n=10 |  | 32.81<br>± 6.79 | 24.06<br>± 3.93 | 25.00<br>± 4.47 | 21.25<br>± 6.67 | 16.56<br>± 5.02 | 19.69<br>± 3.98 | 14.06<br>± 2.52 | 10.63<br>± 3.27 | 28.75<br>± 8.55 | 6.25<br>± 5.02 | 12.50<br>± 5.59 | 2.50<br>± 2.50 | 28.75<br>± 9.33 | 7.50<br>± 5.34 | 10.00<br>± 6.67 | 5.00<br>± 5.00 |
| Gq+CNO<br>: Paired-Paired | n=9 |  | 33.68<br>± 5.36 | 25.35<br>± 6.43 | 25.00<br>± 5.13 | 14.24<br>± 3.54 | 20.14<br>± 4.97 | 25.00<br>± 7.27 | 13.89<br>± 5.24 | 9.38<br>± 4.29 | 2.78<br>± 1.84 | 12.50<br>± 6.59 | 13.89<br>± 7.35 | 11.11<br>± 7.35 | 25.00<br>± 9.32 | 11.11<br>± 7.35 | 0.00<br>± 0.00 | 1.39<br>± 1.39 |

<sup>#</sup>, 2-trial block

Latent inhibition - LHb lesion (supplement figure 1)
| Group |  | Day | Phase 1: Unpaired conditioning |  |  |  |  |  | Phase 2: Paired conditioning |  |  |  |  |  |
| --- | --- | --- | --- | --- | --- | --- | --- | --- | --- | --- | --- | --- | --- | --- |
|  |  |  | 1 | 2 | 3 | 4 | 5 | 6 | 7 | 8 | 9 | 10 | 11 | 12 |
| Sham | n=6 |  | 10.94 | 4.17 | 4.17 | 5.73 | 7.29 | 3.65 | 25.00 | 23.96 | 13.02 | 28.13 | 21.88 | 15.63 |
| : CS alone-Paired |  |  | ± 5.69 | ± 3.09 | ± 2.08 | ± 3.46 | ± 3.84 | ± 2.34 | ± 4.03 | ± 6.39 | ± 4.07 | ± 10.52 | ± 9.20 | ± 6.99 |
| Lesion | n=6 |  | 6.25 | 4.69 | 10.94 | 2.08 | 5.73 | 4.69 | 21.35 | 28.65 | 10.94 | 4.69 | 13.02 | 15.10 |
| : CS alone-Paired |  |  | ± 4.03 | ± 2.65 | ± 5.69 | ± 2.08 | ± 3.17 | ± 1.76 | ± 7.92 | ± 11.56 | ± 4.82 | ± 2.10 | ± 6.11 | ± 6.67 |
| Sham | n=6 |  | - | - | - | - | - | - | 15.63 | 28.65 | 28.65 | 20.31 | 14.06 | 15.63 |
| : Paired alone |  |  |  |  |  |  |  |  | ± 4.35 | ± 9.93 | ± 8.09 | ± 4.55 | ± 4.62 | ± 4.97 |
| Lesion | n=6 |  | - | - | - | - | - | - | 29.69 | 26.04 | 23.44 | 13.02 | 16.67 | 8.33 |
| : Paired alone |  |  |  |  |  |  |  |  | ± 6.93 | ± 7.97 | ± 4.47 | ± 5.67 | ± 2.64 | ± 3.19 |

Retardation - inhibition of LHb with DREADDs (supplement figure 2)
| Group |  | Day | Phase 1: Unpaired conditioning |  |  |  |  |  | Phase 2: Paired conditioning |  |  |  |  |  |
| --- | --- | --- | --- | --- | --- | --- | --- | --- | --- | --- | --- | --- | --- | --- |
|  |  |  | 1 | 2 | 3 | 4 | 5 | 6 | 7 | 8 | 9 | 10 | 11 | 12 |
| Gi+CNO | n=8 |  | 10.94 | 14.84 | 9.38 | 12.11 | 5.86 | 9.38 | 8.98 | 10.55 | 14.06 | 13.28 | 3.52 | 10.16 |
| : Unpaired-Paired |  |  | ± 3.07 | ± 7.08 | ± 4.18 | ± 4.85 | ± 1.72 | ± 3.34 | ± 3.09 | ± 5.31 | ± 6.94 | ± 4.04 | ± 1.72 | ± 3.58 |
| Gi+VEH | n=9 |  | 25.35 | 14.58 | 11.11 | 9.72 | 9.38 | 8.68 | 4.51 | 2.08 | 4.51 | 1.74 | 6.60 | 5.56 |
| : Unpaired-Paired |  |  | ± 5.35 | ± 3.21 | ± 3.65 | ± 3.70 | ± 3.97 | ± 1.71 | ± 1.81 | ± 1.38 | ± 1.39 | ± 1.06 | ± 3.77 | ± 1.63 |
| EGFP+CNO | n=7 |  | 18.30 | 18.30 | 8.48 | 12.05 | 12.95 | 5.36 | 7.59 | 5.36 | 3.57 | 2.68 | 8.93 | 4.02 |
| : Unpaired-Paired |  |  | ± 4.75 | ± 7.76 | ± 4.71 | ± 6.41 | ± 5.08 | ± 3.72 | ± 3.85 | ± 2.53 | ± 1.26 | ± 1.26 | ± 3.51 | ± 1.48 |
| Gi+VEH | n=9 |  | - | - | - | - | - | - | 9.03 | 16.67 | 11.11 | 5.21 | 11.81 | 10.07 |
| : Paired alone |  |  |  |  |  |  |  |  | ± 2.05 | ± 3.53 | ± 3.22 | ± 1.73 | ± 3.45 | ± 3.56 |

